# Factors affecting germination and establishment success of an endemic cactus of the Chihuahuan Desert

**DOI:** 10.1101/2020.12.08.396481

**Authors:** Eder Ortiz-Martínez, Jordan Golubov, María C. Mandujano, Gabriel Arroyo-Cosultchi

**Affiliations:** Departamento El Hombre y su Ambiente. Universidad Autónoma Metropolitana Xochimilco, Calzada del Hueso 1100. Col Villa Quietud, 04960 México. DF., México..; Departamento El Hombre y su Ambiente. Universidad Autonoma Metropolitana Xochimilco, Calzada del Hueso 1100. Col Villa Quietud, 04960 México. DF., México; Departamento de Ecología y Recursos Naturales, Facultad de Ciencias, Universidad Nacional Autónoma de México, México D.F. 04510, México..; Instituto de Ecología, Universidad Nacional Autónoma de México (UNAM), Departamento de Ecología de la Biodiversidad, Laboratorio de Genética y Ecología, Apartado Postal 70-275, Ciudad Universitaria UNAM, 04510 México, DF., México

**Keywords:** Facilitation, seedling recruitment, seed germination, *Cephalocereus polylophus*, seed and seedling limitation, Cactaceae

## Abstract

Seed and seedling are the most critical stages of cacti life cycle. From the thousands of seeds produced in a reproductive season, only a small fraction gets to germinate, the rest gets lost due to predation or gets potentially buried in the seed bank. These early stages depend on facilitation by nurse plants for germination and seedling recruitment. In this paper, we aim to describe some aspects of the recruitment of *Cephalocereus polylophus*. We tested the viability of seeds with different storage times as an indicator of their potential to form a short-term seed bank. Through the analysis of seed germination and seedlings survival under the canopy of two nurse plant species and open areas, we aimed to assess the importance of facilitation for recruitment. A predator exclusion experiment was used to evaluate the intensity of herbivory on seeds and seedlings of different developmental stages. Seeds had germination rates above 90%, even after two years of storage. Seed germination was only registered under one of the two nurses. After two years, up to 19% of the seedlings planted under both nurse plants survived. Protection against herbivores increased survival chances from 30 to 52 % for all age-group seedlings. Considering that facilitation is a crucial interaction for *C. polylophus*, future conservation programs should include the protection of plant communities.

## 1 Introduction

Seedlings are one of the central components of plant population dynamics (Harper, 1974). The recruitment of seedlings, understood as the process by which new individuals are added to a population, includes seed germination, seedling survival, and seedling growth (Eriksson and Ehrlén, 2008). Seedlings play a central role in the conservation of many plant species (Valiente-Banuet and Ezcurra, 1991; Godínez-Álvarez et al., 2003; Pierson et al., 2013); however, it has been hypothesized that populations can be severely seedling limited (Rees, 1994) by biotic and abiotic factors, such as high temperatures (Nobel, 1984; Suzán-Azpiri and Sosa, 2006; Miranda-Jácome et al., 2013), low and unpredictable water availability (Holland and Molina-Freaner, 2012) and strong predation pressures (García-Chávez et al., 2010; Holland and Molina-Freaner, 2012), all of which negatively impact seed germination and seedling survival (Bowers, 1997; Pimienta-Barrios et al., 2002; Godínez-Álvarez et al., 2003; Rojas-Sandoval and Meléndez-Ackerman, 2012; Pierson et al., 2013).

Cacti are among the most ubiquitous plant families of North and South American deserts (Guerrero et al., 2019). Many of the cacti species release a large number of seeds to their populations on an annual basis, but most are rapidly removed by birds, rodents, and ants before having the chance to germinate (Sosa and Fleming, 2002; García-Chávez et al., 2010; Holland and Molina-Freaner, 2012).

Plant facilitation has been documented to be a positive interaction for seedlings, especially during the early stages of establishment, in which benefactor species (nurse) interacts with beneficiary species (facilitated) (Verdú et al., 2010). The importance of this type of association has been reported for several cacti species, nurse plants from the families Fabaceae, Asteraceae, Mimosaceae, and even Cactaceae are known to be crucial for seed germination and seedling survival (Nobel, 1984; Franco and Nobel, 1989; Valiente-Banuet and Ezcurra, 1991; Mandujano et al., 1998, 2001; Rojas-Sandoval and Meléndez-Ackerman, 2012). It has been demonstrated that nurse plants protect seedlings by decreasing the maximum soil surface temperature beneath their canopies (Nobel, 1980; Franco and Nobel, 1989; Miranda-Jácome et al., 2013; Munguía-Rosas and Sosa, 2008), reducing evapotranspiration (Valiente-Banuet et al., 1991), improving soil properties (Montesinos-Navarro et al., 2016; Munguía-Rosas and Sosa, 2008) and protecting seeds and seedlings from predation (Mcauliffe, 1984; Valiente-Banuet and Ezcurra, 1991; Holland and Molina-Freaner, 2012; Sosa and Fleming, 2002).

The conservation of many cacti species would benefit from a clear understanding of the biotic and abiotic factors that play a role in processes that constrain population dynamics. Using *Cephalocereus polylophus* as a study species, we aim to answer the following questions: what is the short term seed viability?, what is the effect of different light intensities on germination and seedling development?, how important is facilitation for seed germination and seedling survival?, how intense is seedling predation and which are the most vulnerable stages of seedling development?

## 2 Materials and methods

### 2.1 Subject species

*Cephalocereus polylophus* (DC.) Britton & Rose (= *Neobuxbaumia polylopha*) is a columnar cactus that occurs naturally in canyon regions covered with deciduous forest and calcareous soils. It is distributed in an area of approximately 6,000 km^2^ constrained to six isolated locations in central Mexico (IUCN, 2019). Populations from the Barranca de Metztitlán Biosphere Reserve are dispersed but some have high densities (0.67 ind/m2 ± 0.28 (Arroyo-Cosultchi et al., 2016). The flowering season is between May and July, flowers are hermaphroditic, large (mean±SD length 4.63 ± 0.08 cm) with a dark pink perianth, anthesis is nocturnal and lasts for one and rarely two nights (Anderson, 2001; Arroyo-Cosultchi et al., 2010). Flowers are nectariferous and are visited by bats and hummingbirds (personal observation) (Cornejo-Latorre et al., 2011). Mean seed production is 760 seeds per fruit (mean seed length and width: 2.68 × 1.85 mm) (Arroyo-Cosultchi et al., 2007), which are dispersed by bats, birds, and ants during July and August (Arroyo-Cosultchi et al., 2016).

### 2.2 Study area

The study was carried out in the Barranca de Metztitán Bio-sphere Reserve (RBM) in the vicinities of the town of San Miguel Almolón (20° 43’ 32.8” N - 98° 54’ 56.9” W), in the state of Hidalgo, Mexico. The RBM is considered one of the most outstanding cactus regions in the country, 70 species are registered in the region, 11.42 % of which are endemic to the site and 15 % of them are under some category of risk (Sánchez-Mejorada, 1978; CONANP, 2003). The climate is dry, semi-warm with summer rains (BS0hw), mean annual temperature is around 20.7 °C (14.1 - 27.3 °C), mean annual precipitation is 388.40 mm (min-max 92 - 799 mm) with about 85 % of annual precipitation occurring during summer and fall (CONAGUA, Meteorological Station San Cristobal, Metztitlán; 20° 38’29” N - 98° 49’ 42.96” W). During the study period (June 2014 – June 2015), mean annual rainfall was 388.40 mm (92-799 mm) with peaks occurring in September. The dominant vegetation type is xerophytic scrub (CONANP, 2003). The vegetation is classified as deciduous forests and microphyllous scrubs, dominated by the columnar cactus *Cephalocereus senilis* (Haw.) Pfeiff., Allg, and *Isolatocereus dumortieri* (Scheidw.) Backeb. (Cruz and Pavón, 2013) as well as shrubby legumes (Herce et al., 2013).

Three mature fruits from 30 different *C. polylophus* individuals were collected, during the months of June and July of 2012, 2013 and 2014. Immediately after collecting the fruits, seeds were cleaned, dried and stored in paper bags at room temperature to avoid fungal infestation.

### 2.3 Seed limitation

#### 2.3.1 Seed viability

Seed viability of *C. polylophus* over time was assessed in September 2014. Seeds collected in 2012 (twenty-seven months old), 2013 (fifteen months old), and 2014 (three months old) were germinated under lab conditions. Three hundred seeds of each year (900 total seeds) were sown in Petri dishes with 1 % bacteriological agar. Thirty replicates (10 seeds per petri dish) per year, were placed in an environmental chamber under controlled temperature and photoperiod (Lab-Line Biotronette 845, 28-30 °C and 12 h photoperiod). Germination was registered daily for 30 days. A GLM with a binomial error distribution was used to determine differences in germination between seed age. The height and diameter of each seedling were measured as a proxy of seedling vigor (*n*=830) at the end of the experiment.

#### 2.3.2 Seed germination under different PAR intensities

To assess how germination is affected by different light intensities, seeds collected in 2013 were subject to four different light treatments under controlled laboratory conditions. Seeds were germinated inside an environmental chamber (Lab Line Biotronete 845), calibrated to perform at a maximum photosynthetically active radiation (PAR) of 394 *μ*mol ^−2^ s^−1^, 28-30 °C and 12h photoperiod). The full radiation of the environmental chamber was used as the 100 % PAR treatment. Greenhouse shade cloth was used to simulate different PAR treatments as follows: 45 % (182 *μ*mol m^−2^ s^−1^), 25 % (103 *μ*mol m^−2^ s^−1^) and 10 % (38 *μ*mol m^−2^ s^−1^), PAR was measured with a Li250A LI-COR sensor. Every treatment (*n* = 4 PAR intensities) had 30 replicates, each with 20 seeds sown in Petri dishes with 1 % bacteriological agar, for a total of 120 experimental units. Germination was registered daily for 30 days. In order to assess the effect of different PAR exposition on seedling development, the length and diameter of 100 seedlings from every treatment were measured with a digital caliper to the nearest 0.01 (*n* = 400 seedlings). A GLM with a binomial error distribution was used to test differences in germination between light treatments and ANOVA for the length and diameter response variables followed by Tukey *post hoc* tests.

#### 2.3.3 Germination and seed survival in habitat

The germination and predation of *Cephalocereus polylophus* seeds in habitat were evaluated during the 2014 rainy season (August). Experimental units for both experiment consisted of ten seeds glued with liquid silicone to a thin plastic mesh. A pilot experiment with this method showed that the liquid silicone did not interfere with germination (E. Ortiz-Martínez unpublished data).

##### Nurse effect

To assess if deciduous shrubs facilitate *C. polylophus* germination, we tested the germination of seeds beneath the canopy of two shrub species, *Croton mazapensis* Lundell and *Hoverdenia speciosa* Nees as well as in open areas. Two experimental units were placed beneath individuals of each nurse species and in open areas (three treatments, 23 individuals per treatment *n* =1,380 seeds). The experimental units were fixed with nails to the soil to avoid its removal by wind or water and to keep seeds in contact with the soil. The number of germinated seeds was counted 30 days after sowing.

##### Seed predation

To determine if nurse plants protect *C. polylophus* seeds from predation, we tested seed removal beneath the canopy of *C. mazapensis*, *H. speciosa* and in open areas. The seed predation experimental design consisted of twenty 0.5 *m*^2^ plots located randomly in a bushy area. In ten plots, three experimental units were placed inside 1 mm aperture wire mesh boxes (10 cm^3^) which functioned as protection against small vertebrate granivores. The other 10 plots also had tree experimental units, but these were left unprotected. Seed germination and the number of removed seeds were counted 30 days after sowing (*n* =600 seeds).

### 2.4 Seedling limitation

#### Nurse effect

The importance of nurse plants for the seedling establishment was evaluated from 2014 to 2016, we tested the survival of *C. polylophus* seedlings under the canopy of *C. mazapensis*, *H. speciosa* and in open spaces. Seeds were germinated in laboratory ten months before they were transplanted in habitat. A month before the beginning of the experiment, seedlings were hardened for a month in a greenhouse. Experimental units consisted of 12 cm biodegradable jiffy pots filled with soil from the study site and ten 10-month-old seedlings transplanted in them. Twenty-three experimental units were planted under the canopy of 23 different shrubs of each species and 23 open areas. Seedling survival was recorded daily for the first 12 days and then 1, 2, 5, 12, and 24 months after their introduction to habitat.

#### Seedling predation

The predation of the seedlings of different age was measure was evaluated from 2014 to 2016. Seeds were germinated in laboratory five, ten, and twelve months before they were transplanted in habitat. Experimental units consisted of 12 cm biodegradable jiffy pots filled with study area soil, and 10 seedlings of each one of the age groups transplanted in them. The experimental units were transplanted in twenty 0.5 *m*^2^ plots located randomly in partially shaded sites. In 10 plots, the experimental units with 5, 10 and 12 month-old seedlings were protected by a 1 mm aperture wire mesh boxes (10 cm^3^). In the other 10 plots, the experimental units were left unprotected. Seedling survival was recorded daily for the first 12 days and then 1, 2, 5, 12, and 24 months after introduction.

Differences among treatments for seedling survival were analyzed using GLM with binomial error distribution and logit as a link function, and orthogonal contrasts. Seedlings survival curves from the age and exclusion experiments were analyzed using the Kaplan-Meir method, the statistical differences among treatments were tested using the Log-Rank test. Analysis were performed with survival and survminer packages (Therneau, 2020; Kassambara et al., 2020) in R (R Development Core Team, 2020) v.4.0.2.

## 3 Results

### 3.1 Seed viability

Germination rate of *C. polylophus* seeds under laboratory conditions was very high (95.7 ± 5.9 %); there were no significant differences among the seeds of different age (*χ*^2^ = 0.3496, gl = 2, P = 0.8396). Seedlings length (F_2,826_ = 455, P < 0.001) and diameter (F_2,826_ = 235, P< 0.001) were statistically different, seedlings from 2014 were the largest (Table 1). Correlation between seedling length and diameter was weak but significant (r = −0.160, P < 0.01).

**Table 1.** Germination rate, length and diameter (mean ± standard error) of 30 days old *Cephalocereus polylophus* seedlings from different age seeds. Different letters indicate significant differences (*P* < 0.01).

| Year of collection | Germination rate (%) | Length (mm) | Diameter (mm) |
| --- | --- | --- | --- |
| 2012 | 93.66 a | 8.15±0.05 c | 3.92±0.03 b |
| 2013 | 98.33 a | 8.48±0.05 b | 4.18±0.03 a |
| 2014 | 95.66 a | 10.37±0.07 a | 3.54±0.02 c |

### 3.2 Seed germination under different PAR intensities

There were no significant differences in germination under different Photosynthetically Active Radiation (PAR) intensities (*χ*^2^ = 3.12, gl = 3, P = 0.3732). All treatments had germination rates higher than 89 %. Significant differences were found in the length of seedlings exposed to different PAR intensities (F_3,144_ = 123.29, P < 0.05). Seedlings were longest at the lower PAR intensities (Figure 1 a). Seedlings diameter was also significantly different (F_3,144_ = 69.95, P < 0.05), but Tukey Test showed no statistical differences between the seedlings exposed to the lowest and highest PAR intensities (Figure 1 b). Correlation between seedling length and diameter was weak but significant (r = 0.279, P < 0.05).

**Fig. 1.**
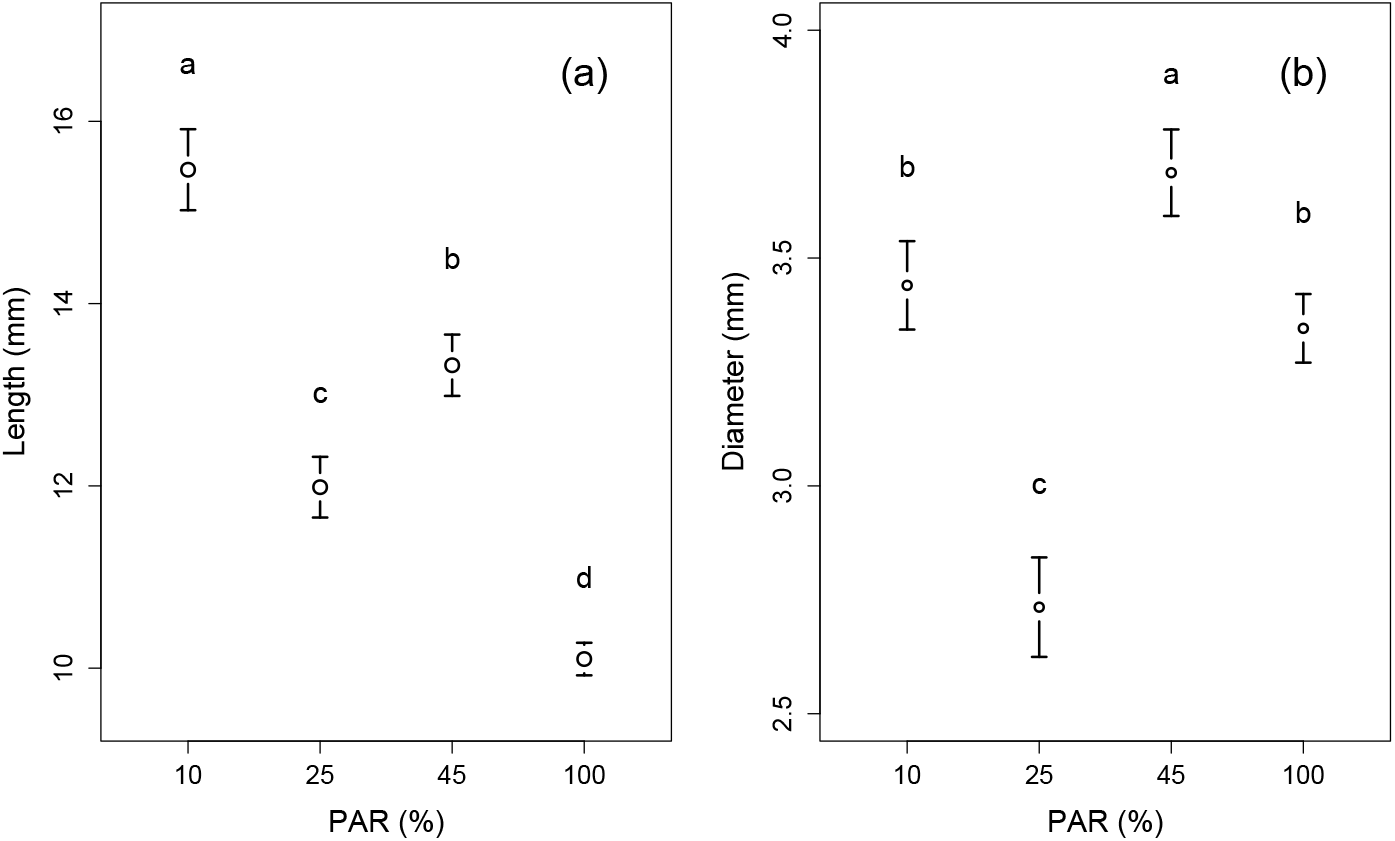
Length and diameter (Mean ± confidence interval 95 %) of *Cephalocereus polylophus* seedlings germinated under different Photosynthetically Active Radiation (PAR) intensities. Different letters indicate differences among PAR treatments for the response variable (*P*< 0.05).

### 3.3 Germination and seed survival in habitat

After thirty days, only 2.39 % of the seeds placed under *Croton mazapensis* germinated. No germination was recorded under the canopy of *Hoverdenia speciosa* nor in open spaces. During the same period, 65.0 % and 67.6 % of the seeds placed under *C. mazapensis* and *H. speciosa* were removed or predated, while in open spaces 76.9 % of the seeds had the same fate. In the exclusion experiments, 6.6 % of the protected seeds and 62.66 % of the exposed seeds were removed during the first 12 days. Exclusion cages only avoided the removal of seeds by vertebrate granivores. Ants from the species *Campanotus planatus* Roger were observed removing seeds of *C. polylophus*.

### 3.4 Seedlings survival in habitat

Nurse plant protection was crucial for seedling survival. While every seedling in open areas died during the first month, 19.1 % of the seedlings transplanted under *H. speciosa* and 18.7 % of those under *C. mazapensis* survived after 24 months (Figure 2).

**Fig. 2.**
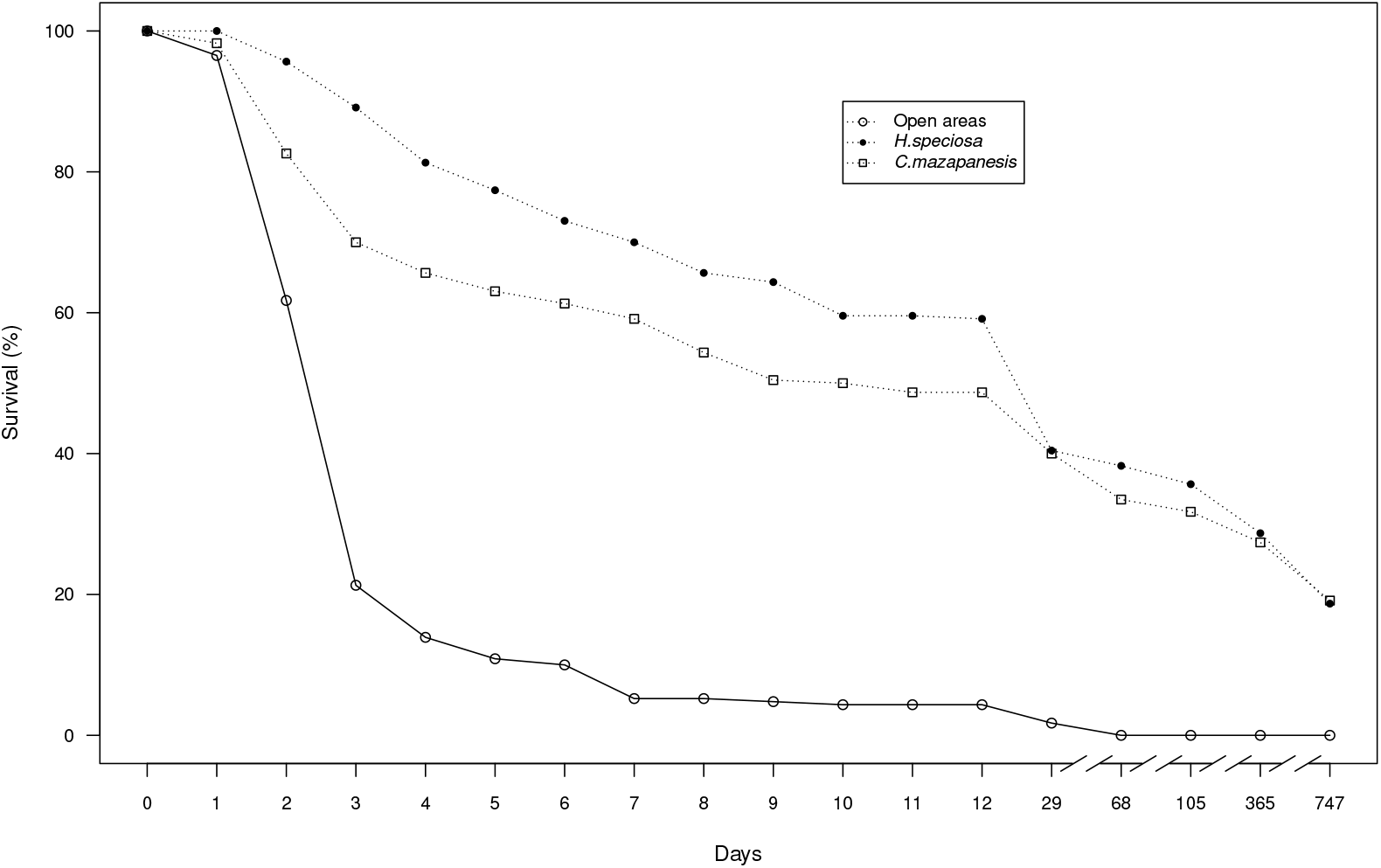
Survival of 10 month-old *Cephalocereus polylophus* seedlings transplanted under two nurse species and open spaces. *n_initial_*=230 seedlings/treatment.

The age of the seedlings and the protection against predators were determinant for seedling survival (*χ*^2^ = 250.08, gl = 5, P < 0.05). Analyses of the survivorship curves indicated significant differences among treatments (Long-Rank test P<0.0001). No significant differences were detected between the protected ten month-old seedlings and exposed twelve-month-old seedlings (P<0.4463). By the end of the experiment, survival was higher for the oldest seedlings, regardless of whether they were protected or exposed (95 % of the protected and 65 % of the exposed twelve-month old seedlings survived after 24 months). Protection against herbivores increased survival chances from 30 to 52 % for all age-group seedlings (Figure 3).

**Fig. 3.**
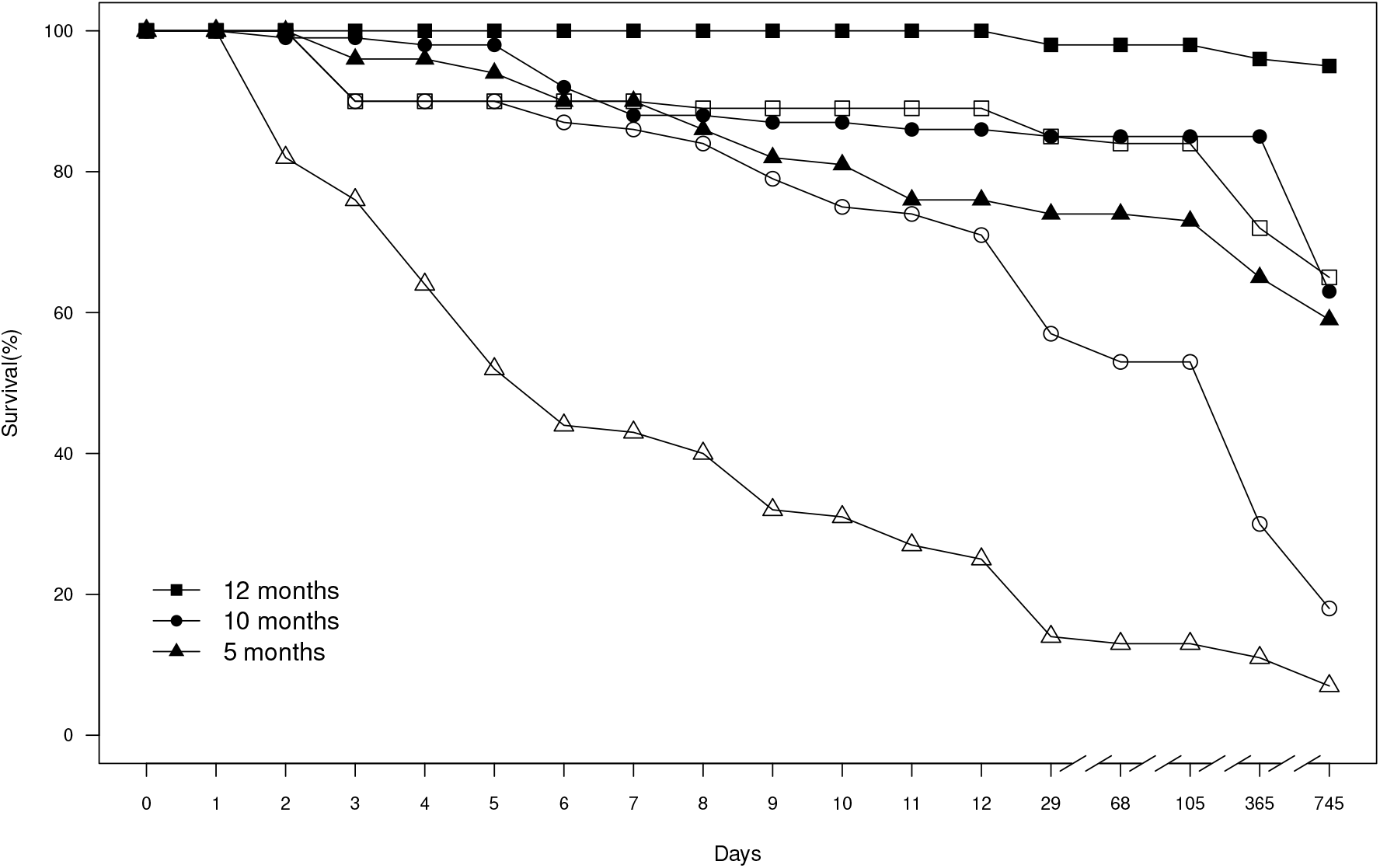
Survival of *Cephalocereus polylophus* seedlings of different age and exclusion treatment after 24 months. Treatments with exclusion=symbol in bold and without exclusion=symbol without filling. *n _initial_*=100 seedlings/treatment.

## 4 Discussion

Seeds and seedlings are well known to be vulnerable and are usually considered a severe bottleneck to population growth (Harper, 1977; Clark et al., 2007). The low number of seedlings observed in natural cacti populations can be partially explained by post-dispersal constrains which include, seed viability and senescence (Mandujano et al., 1998; Clark et al., 2007). Seed viability varies strongly among cacti species (Rojas-Aréchiga et al., 2013) and senescence has been shown to affect viability in many species (Álvarez-Espino et al., 2014; Lindow-López et al., 2018). In the case of *Cephalocereus polylophus*, seed viability does not seem to be a relevant component as germination rates were high even considering senescence, at least during the studied period, which suggests that the species is not limited by seed viability and could even potentially form a short-term persistent seed bank (Thompson et al., 1993; Ordoñez, 2016).

Abiotic components have a strong influence in the seed-seedling interphase, especially in arid and semi-arid environments that pose varying and extreme abiotic conditions (Eriksson and Ehrlén, 2008; Miranda-Jácome et al., 2013). Seeds in these environments must balance between an excess of light that can lead to extreme temperatures found in open spaces and the competition for light under the canopy of other plants (Landero and Valiente-Banuet, 2010). The need for light for germination in the Cactaceae has been shown to be phylogenetically fixed for some groups (Rojas-Aréchiga et al., 2013). Although the intensity of the Photosynthetically Active Radiation (PAR) did not affect the germination of *C. polylophus* seeds under laboratory conditions, as with other species of the tribe Cacteae (Rojas-Aréchiga et al., 2013), the intensity of light does seem to be important for seedling development. Seedlings germinated in the laboratory under low PAR exposure were longer than those exposed to high PAR. In natural conditions the seedlings of columnar cacti such as *Stenocereus thurberi* (Nolasco, 1997) and *C. mezcalaensis* (Landero and Valiente-Banuet, 2010) grow faster in shaded places such as that found under nurse plants.

The microclimates provided by nurse plants reduce the adverse effects of extreme environmental variables and increase the chances of survival of facilitated plants (Turner et al., 1966; Valiente-Banuet and Ezcurra, 1991; Méndez et al., 2006). *Cephalocereus polylophus* was particularly sensitive to the exposure to the high radiation and temperatures in its habitat, indicating that facilitation is a crucial association for the species. The high seedling mortality in open spaces during the first month is consistent with the pattern reported for other species with CAM metabolism (Turner et al., 1966; Jordan and Nobel, 1979; Valiente-Banuet and Ezcurra, 1991) and other species of *Cephalocereus* (Landero and Valiente-Banuet, 2010).

The rapid removal of seeds from the soil reduces the number of seeds available for germination and has been widely documented for several columnar cacti species (Turner et al., 1966; Valiente-Banuet and Ezcurra, 1991; Méndez et al., 2006; Miranda-Jácome et al., 2013). Perennial plants from semiarid environments experience intense granivory (Méndez et al., 2006; Landero and Valiente-Banuet, 2010), which can reduce the number of seeds available to germinate to only 5 % of the total seeds released in every reproductive season (Turner et al., 1966; Valiente-Banuet and Ezcurra, 1991). In the case of *C. polylophus*, only 2.39 % of seeds were able to escape predation. This is a common phenomenon in Cactaceae, for example 60 to 95 % of *Opuntia rastrera* seeds are removed during the first months after fruiting, mainly by rodents, birds and ants (Mandujano et al., 2001; Montiel and Montaña, 2003), as well as 78 to 100 % of the seeds of *Cephalocereus mezcalaensis* (Landero and Valiente-Banuet, 2010), and 99% of the seeds of *Ferocactus wislizeni* (Bowers, 2000).

After germination, seedlings are also subject to intense predation (Clark et al., 2007). Herbivory is a factor that negatively impacts many species of cacti, specially during the earlier stages of their life cycle (Landero and Valiente-Banuet, 2010; Mandujano et al., 1998; Turner et al., 1966; Mcauliffe, 1984). Seedlings of *C. polylophus* are susceptible to be used as a resource by small grazers, mainly ants, birds, and small mammals, which have been reported to commonly exploit cacti seeds (Méndez et al., 2006; Landero and Valiente-Banuet, 2010). We observed a strong predation on the 5 month-old seedling while most of the twelve months-old seedlings survive after two years, it is clear that the age and the increment of size allow seedlings to escape from the most critical stages of their life cycle (Steenbergh and Lowe, 1977; Valiente-Banuet and Ezcurra, 1991; Munguía-Rosas and Sosa, 2008). Additionally to the reduction of susceptibility to predation, larger seedlings have also larger water storage volume, which provides a buffer against extended droughts (Jordan and Nobel, 1981).

The amount of seeds produced by *C. polylophus* and their high viability suggest a high potential for recruitment (no evident seed limitation), however, seedling establishment is significantly impacted by the removal of the seeds, the dependence on nurse plants, and the high mortality rate of seedlings by both abiotic and biotic factors. Habitat conservation is crucial for nurse-dependent species such as *C. polylophus* since they depend largely on the size, density and complexity of their nurse species (Bruno et al., 2003). Nurse plant association and the reintroduction of mature seedlings should be the main method in future restoration and conservation strategies for *C. polylophus* and other cacti species.

## Acknowledgements

This research is part of the bachelor’s degree studies of Ortiz-Martínez Eder (UAM-X) and the PhD studies of Arroyo-Cosultchi Gabriel. Financial support was provided by CONACyT(165908) to MCM, CONACyT sabbatical leave scholarship to JG and PASPA-DGAPA sabbatical scholarship to MCM. We appreciate the suggestions provided by Mariana Rojas-Aréchiga and Susana Guillén Rodríguez on an earlier version of the manuscript.

## Conflict of interest

The authors declare that they have no conflict of interest.

## Notes

### Competing Interest Statement

The authors have declared no competing interest.

